# Conscious perception and the role of the basal ganglia: preliminary findings from a deep brain stimulation study

**DOI:** 10.1101/2022.11.15.516581

**Authors:** E.A. Boonstra, M.N. Bais, M.R. van Schouwenburg, P. van den Munckhof, D.J.A. Smit, D. Denys, H.A. Slagter

## Abstract

Conscious perception is thought to depend on global amplification of sensory input. In recent years, the basal ganglia have been implicated in gating conscious access due to their consistent involvement in thalamocortical loops. However, much of the evidence implicating the basal ganglia in these processes in humans is correlational. The current study is a preliminary investigation in four patients to explore whether deep brain stimulation (DBS) in the basal ganglia might improve conscious perception. In our study, treatment-resistant obsessive-compulsive disorder (OCD) patients with a striatal DBS implant completed two canonical conscious perception tasks: emotion-induced blindness and backward masking. We found preliminary evidence in support of a role played by the basal ganglia in conscious perception at the behavioral level: patients performed better when stimulation was active, but we could not establish neural effects corresponding to these behavioral findings, possibly due to our small sample size. We discuss the potential implications and limitations of our study and delineate avenues for future research.

## Introduction

The relationship between consciousness and the brain is often considered as one of the major frontiers of contemporary science. Within the context of conscious visual perception, one major outstanding question concerns how a stimulus becomes accessible for report. Influential theoretical accounts have proposed that such accessibility requires “broadcasting” of a stimulus to the whole brain through thalamocortical circuits (Crick & Koch, 2003; Dehaene & Changeux, 2011; Edelman, 2003). The basal ganglia (BG) are a cluster of subcortical nuclei located deep in the brain, which modulate these circuits (Smith et al., 2004), and as such, may gate conscious perception. For decades we have known that BG structures contribute to sensory and perceptual processes (Afrasiabi et al., 2021; Alexander & Crutcher, 1990; Arsalidou et al., 2013; Brown et al., 1997; Seger, 2013), in line with the notion that the BG not only perform action selection, but fulfill a general selection function, including selection for conscious perception (e.g., Redgrave et al., 1999). It is notable in this respect that fMRI studies consistently show differences in BOLD activity in the striatum between consciously perceived and unconscious stimuli using backward masking tasks (Bisenius et al. 2015). Another repeated finding is a relationship between striatal D2 receptor binding and performance on conscious perception tasks (Bisenius et al., 2015; Slagter et al., 2010, 2012; Van Opstal et al., 2014; but see Boonstra et al., 2020). Most of these studies employ neuroimaging, and thereby lack temporal precision and/or a direct measure of BG activity. This has left it largely unclear how the BG may contribute to conscious perception in humans.

In recent years, deep brain stimulation (DBS) electrodes have proven an effective tool to not only modulate BG structures, but also to measure their activity. DBS involves an invasive procedure leveraged to alleviate symptoms of several treatment-resistant neurological and psychiatric pathologies such as Parkinson’s disease (Almeida et al., 2017), anorexia nervosa (Whiting et al., 2018), depression (Dandekar et al., 2018), and obsessive-compulsive disorder (OCD, Baldermann et al., 2021). A previous study employed an attentional blink task to probe conscious perception in patients with DBS electrodes implanted in the ventral striatum (Slagter et al., 2017). The attentional blink comprises a deficit in consciously perceiving the second of two targets (T1 and T2) whenever it follows the first target within 100–500 ms in a rapid stream of distractors (Raymond et al., 1992). Patients performed this task while intracranial EEG (iEEG) activity from the ventral striatum was recorded using the DBS electrodes without stimulation. Intracranial EEG recordings revealed a consistent burst in theta-band activity only when T2 was consciously perceived (Slagter et al., 2017). Moreover, increased activity in α and low β frequencies after perceiving T1 was observed in trials in which subjects subsequently failed to perceive T2, possibly reflecting attentional capture by T1. These findings support the notion that the BG contribute to conscious experience. Yet, like neuroimaging, they only provide correlational evidence.

In the present study, we present preliminary data from a small sample of four treatment-resistant OCD patients with DBS electrodes implanted in the ventral striatum. The efficacy of DBS treatment in OCD patients is still under active investigation, but evidence thus far encourages further study (Acevedo et al., 2021; Graat et al., 2021; van Wingen et al., 2022; Wu et al., 2021). Our study goes beyond past work in that we were able to record iEEG during DBS. We were thereby able to causally manipulate and measure striatal activity simultaneously, while patients performed two canonical conscious perception paradigms: emotion-induced blindness (EIB) and backward masking (BM). This setup allowed us to establish a more conclusive role for the ventral striatum in conscious perception compared to the earlier study by our group. We concurrently measured scalp EEG, permitting investigation of the effects of DBS on common event-related potentials (ERPs) markers of conscious perception (see Hypotheses below). Moreover, the two tasks allowed us to probe a participant’s stimulus detection ability both when the perception of a target is complicated by an emotionally valenced (positive or negative; EIB), as well as a neutral disturbance (BM). Taken together, this setup allowed us to examine if causal perturbation of striatal activity would affect conscious perception within the context of these tasks.

### Hypotheses

Emotion-induced blindness refers to an impaired awareness of stimuli appearing soon after an irrelevant, emotionally arousing stimulus (Most et al., 2005). It is therefore an effect similar to the attentional blink, except that instead of T1, emotionally arousing or valence neutral distractors are presented 200-800 ms before the target. This manipulation of emotional arousal allowed us to vary the degree to which distractors may capture attention in an attempt to extend previous findings (Slagter et al., 2017).

Behaviorally, OFF DBS, we expected to find an effect akin to a standard attentional blink, based on past EIB studies (Kennedy et al., 2014; Most et al., 2005), as reflected in lower accuracy when the lag between the distractor and target is short rather than long. We expected this difference to be larger for targets preceded by negative distractors compared to neutral distractors due to increased attentional capture. In terms of ERPs derived from scalp EEG data, a usual finding for the EIB task is that N2 and P3 components are suppressed for targets following an emotionally arousing distractor (Kennedy et al., 2014). We expected to replicate these findings for targets following a negative distractor OFF DBS.

In the backward masking task, the order of distractor and target is reversed. A target is presented first, followed by a mask (akin to a distractor) at a variable delay, known as the stimulus onset asynchrony (SOA). In this task, patients indicated whether the target, a number, was smaller or larger than five. Commonly, accuracy is better when the delay between target and mask is long rather than short (Breitmeyer, 2007). Behaviorally, OFF DBS, we expected to replicate this finding in terms of improved accuracy for longer SOAs. Our task departs from the standard version in that patients do not only report on the target, but simultaneously indicate the confidence in their response (sure/unsure). Like accuracy, we expected response confidence to improve with increased SOA duration. The reason to include a confidence measure is that the ventral striatum is thought to be involved in determining self-confidence (Kiverstein et al., 2019). We expected DBS ON to improve patients’ accuracy and response confidence. In terms of scalp ERPs, increased ERP amplitude is usually found as SOA increases (Del Cul et al., 2007).

Our study is novel in that we could investigate changes in neural oscillatory activity recorded from the ventral striatum within the context of an EIB and BM task ON and OFF DBS. Based on our previous iEEG attentional blink study (Slagter et al., 2017), OFF DBS, we expected that only consciously perceived targets would be associated with an increase in *θ* activity between 150 and 400 ms after stimulus onset. In the EIB task, we also expected that failure to perceive the target would be foreshadowed by a distractor-induced increase in *α* and low *β* oscillatory activity. Moreover, in the case of EIB, we expected DBS ON to reduce capture by negative distractors and thereby to improve accuracy. At the neural level, we expected this change to be reflected in a reduced activity in *α* and early *β* frequencies immediately following the distractor, as well as an increased *θ* burst and larger N2 and P3 amplitudes to seen targets. In the BM task, we similarly expected the stimulation to increase induced *θ* activity and N2 and P3 amplitudes to seen targets.

Below, we present preliminary findings against the background of earlier work suggesting that DBS may alter patient’s capacity for conscious perception. However, given our small sample size of only four patients, we present solely descriptive statistics and graphical analysis, necessitating more research to substantiate these findings.

## Methods

### Participants

Eleven therapy-resistant OCD patients eligible for deep brain stimulation (DBS) were recruited from the outpatient clinic for DBS at the department of Psychiatry of the Amsterdam University Medical Center, location Academic Medical Center (AMC) from 2015-2020. Alcohol or substance abuse in the past 6 months was a reason for exclusion from the study. Out of the eleven patients recruited, four participated in the tasks comprising the present study (mean age: 48 years, range: 35-61, 3 women, 1 man). Not all eleven patients completed the tasks of the current study because initially the study protocol spanned a full year with the current tasks at the end, for which it turned out the battery life of the stimulator was not suited. One patient performed at chance level in the backward masking (BM) task and was thus excluded from further analyses, resulting in a sample of three patients for this task (mean age: 43 years, range: 35-54, 3 women). All participants were right-handed and took their standard medications. Medications included clomipramine (37.5-75 mg), quetiapine (300 mg), oxazepam (10 mg), fluvoxamine (25-100 mg), fluoxetine (60 mg), lithium carbonate (1000 mg), mirtazapine (45 mg), olanzapine (15 mg), promethazine (25 mg), and lorazepam (1.5 mg). The study was approved by the medical ethics board of the AMC and all patients provided written consent to participate. The trial was registered in the Netherlands Trial Register (Trial NL7486). Patients also performed several other tasks on different study visits (data to be published elsewhere, e.g., Fridgeirsson et al., 2021).

### Experimental Paradigms

Stimuli were presented at a distance of 70 cm on a 15.4-inch laptop screen (HP 6730b, refresh rate = 60 Hz, resolution 1024×768). Due to limited storage capacity of the DBS recording device, and to maximize the number of trials, patients completed a variable number of trials on each block for both tasks.

#### Emotion-Induced Blindness

The emotion-induced blindness task consisted of a central rapid serial visual presentation of briefly (100 ms) presented pictures of landscapes (6.5° wide and 4.9° high) against a black background. Participants were instructed to identify target in this stream (a rotated landscape 90°), and to indicate whether this target was rotated left or right (Fig. 2). As a third response option they could indicate no rotated picture was present (no target). Before the presentation of a target, which was present in each trial, on approximately 86% of trials, a task-irrelevant distractor image was shown, which could be either a neutral image or a negative image, or on approximately 14% of trials, there was no distractor at all. The valence and arousal of these images were rated by the participants themselves on a scale from 1 to 9 on a separate occasion. The 40 most negatively rated pictures served as negative distractors. As neutral distractors we selected 40 pictures rated both least arousing and least positive (but not negative). The target followed the distractor either after 200 ms or 800 ms. Each trial started with an intertrial interval of 200 ms, after which a stimulus stream consisting of 17 stimuli began. A white central fixation cross was visible throughout the experiment. All stimuli were presented using Presentation (version 14.5; Neurobehavioural Systems) at the center of the screen.

#### Backward Masking

The backward masking task was adapted from Van Opstal and colleagues (2014; see Fig. 1). Participants indicated whether briefly presented masked digits (1, 4, 6, or 9) were smaller or larger than 5 and simultaneously rated their confidence in their response, resulting in four response options (>5 sure, >5 unsure, <5 sure, <5 unsure). Responses were counterbalanced in terms of left and right across participants. Each trial started with the presentation of a central fixation cross (53 point Courier New), which increased in size (106 point Courier New, 150 ms duration), cueing the impending target. The target stimulus (53 point Courier New) then appeared for 16 ms at one of two positions centered at the vertical midline (top or bottom, 3.9° from fixation). Both stimulus locations were equally probable. A mask followed the target (200 ms duration) at a variable stimulus onset asynchrony (SOA), with a duration of 16, 33, 50, 66, or 100 ms. By varying the delay between cue and target dependent on SOA, the delay between cue and mask was held constant at 800 ms. The mask (53 point Courier New) was composed of two letters “E” and two letters “M”, tightly surrounding the target location without superimposing or touching it. All stimuli were black and presented on a white background, using the Psychophysics toolbox for MATLAB (Brainard, 1997). The central fixation cross was visible throughout the experiment.

**Figure 1.**
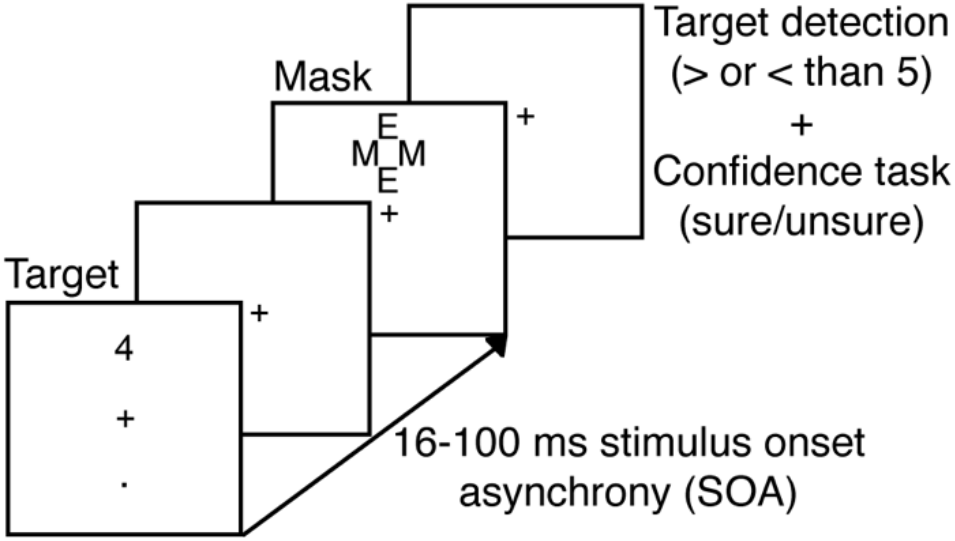
Experimental procedure for one trial of the backward masking task (adapted from Van Opstal et al. 2014)

**Figure 2.**
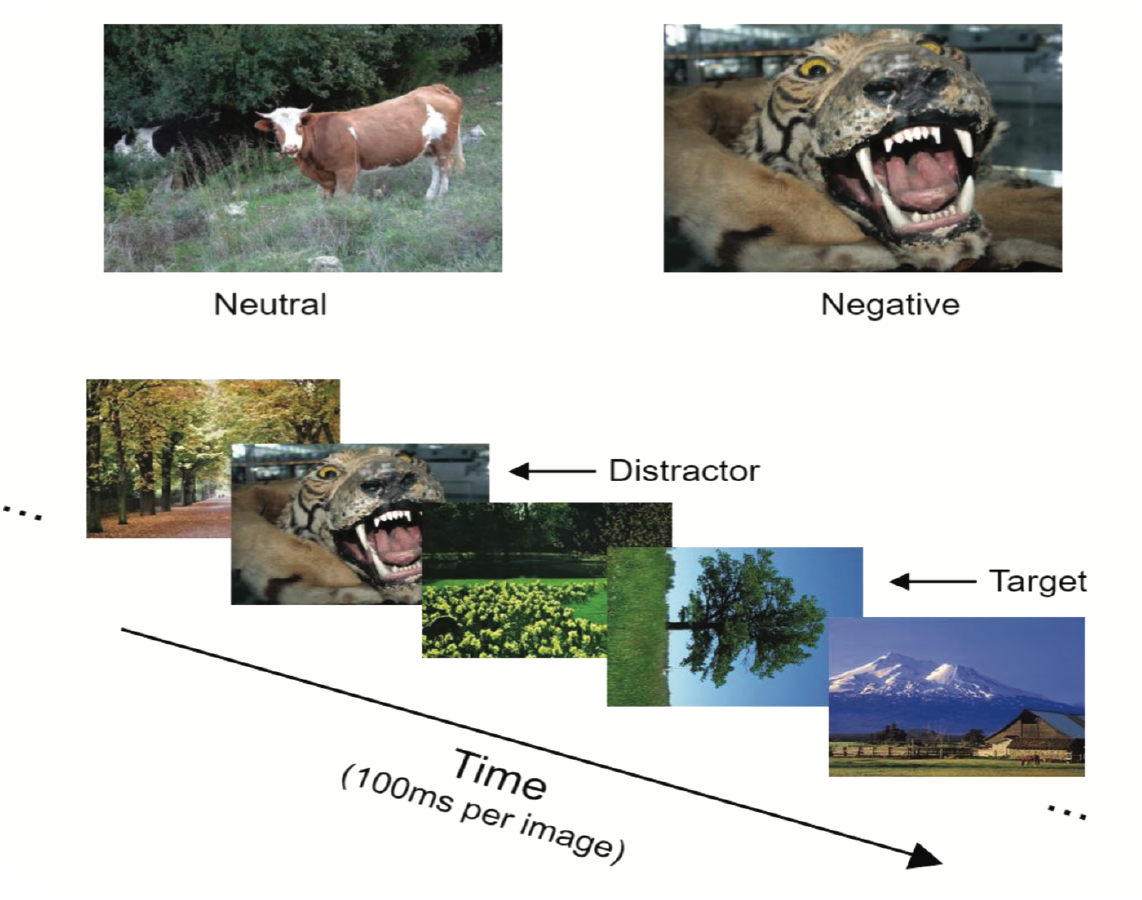
Experimental procedure for one trial of the emotion-induced blindness task

### Neural data recording and preprocessing

Preprocessing was done using the Fieldtrip toolbox (Oostenveld et al., 2011) for Matlab (The MathWorks, Inc. Natick, MA, USA) using custom-written scripts.

#### DBS and Intracranial EEG

Two quadripolar electrodes (Model 3389 Medtronics) with 4 contact points of 1.5 mm each separated by 0.5 mm were implanted stereotactically. Implantation was performed bilaterally through the anterior limb of the internal capsule with the deepest, i.e. most ventral, contact point located in the nucleus accumbens, 3 mm anterior to the anterior commissure and the 3 upper contact points positioned in the ventral part of the anterior limb of the internal capsule (vALIC). The neurostimulator used was the Activa PC + S (Medtronics; model 37604) which allows both for stimulating and recording local field potentials (LFPs) with the recording and stimulation part of the device being completely separated. Stimulation was performed through the middle two contact points at 130 Hz.

Intracranial EEG was recorded from the deepest contact point in the nucleus accumbens using the 8180 Sensing Programmer SW (Medtronic Inc.) with a sampling rate of 422 Hz Using hardware 0.5 Hz high pas filtering and 100 Hz low pass filtering. LFP data were high-pass “firws” filtered (default settings) at 0.5 Hz using a Kaiser window (Widmann et al., 2015). The continuous data were subsequently epoched from −1.5 to 2 s around stimulus presentation to facilitate time-frequency decomposition and baseline corrected to the average activity between −500 ms and −200 ms pre-stimulus in line with Slagter and colleagues (2017). Epochs containing eye blinks surrounding stimulus presentation were rejected based on the visual inspection of the scalp EEG data, that was concurrently measured.

#### Scalp EEG

EEG was recorded at a sampling rate of 1000 Hz using a 64-channel recording system with shielded Ag/AgCl electrodes (ANT Neuro B.V.) following the international 10–10 system. EEG data were high-pass “firws” filtered (default settings) at 0.1 Hz using a Kaiser window (Widmann et al. 2015) and low-pass filtered at 50 Hz due to noise from the DBS implant. Data were offline referenced to the average of all electrodes except LFP, EOG, and mastoid electrodes. The continuous data were subsequently epoched from −1.5 to 1.5 s around stimulus presentation and baseline corrected to the average activity between −500 ms and 200 ms pre-stimulus. We used this time window for both tasks to exclude the distractor in our baseline window in the EIB task, and to remain consistent with iEEG time-frequency analyses. Epochs containing EMG artifacts or eye blinks surrounding stimulus presentation were rejected based on visual inspection. Remaining eye blink artifacts were removed by decomposing the EEG data into independent sources of brain activity using independent component analysis, and removing eye blink components from the data for each subject individually. Epochs were low-pass filtered at 30 Hz for visualization purposes.

### Data analysis

The small sample size precluded the use of inferential statistics; therefore, data were analyzed using descriptive statistics and graphic analysis.

#### Emotion-induced blindness

Three out of four patients completed the task OFF DBS first. Our dependent measure is an accuracy measure reflecting the percentage of correctly identified picture orientation out of all trials (including trials where patients reported no target). Due to our small sample size, relatively low signal-to-noise due to overlapping responses to the RSVP stimuli, and the absence of catch trials, we were unable to compute standard ERPs for this task and only report results of iEEG time-frequency analyses.

#### Backward masking

Two out of three patients completed the BM task OFF DBS first. Our two dependent measures were accuracy (correctly indicating whether the presented number was smaller or larger than 5) and confidence (whether patients were sure or unsure about their response).

To extract event-related potential (ERP) markers of information-processing, we epoched the ERP data to the onset of the mask Del Cul et al (2007). Next, we subtracted the data from the mask-only SOA condition from all other SOA conditions. Finally, we shifted ERP onset back to target onset, to compute target-locked ERPs. Due to the high levels of noise in the data, we focused solely on graphical analysis of the P3b over central parietal scalp sites (P1, Pz, P2, PO3, POz, PO4). This scalp site was used to determine the peak amplitude of this component by averaging the amplitude values 15 ms around the peak sample between 200-400 ms post-target.

## Results

### Emotion-induced blindness

#### Behavior

As shown in Fig. 3, as typically found (Shapiro et al., 1997), patients showcased a robust blink: they less often detected the target on Lag2 (M: 59.4%, SD: 6.6%) compared to Lag8 (M: 68.4%, SD: 10%) distractor-present trials. Moreover, the blink was larger after a negative versus a neutral distractor. That is, within Lag8 trials, patients performed better on average when presented with a negative (M: 70.8%, SD: 11.5%) compared to a neutral distractor (M: 65.9%, SD: 10.4%), while this seeming difference was smaller for Lag2 trials (NEG-M: 60.5%, SD: 7.5%; NEUT-M: 58.4%, SD: 7.9%), resulting in a numerically deeper blink (Lag8-Lag2) after a negative vs. neutral distractor: the emotion-induced blindness effect. Notably, in line with our prediction that DBS would reduce the distractor-induced blink, patients showcased improved accuracy when DBS was ON (M: 63.3%, SD: 6.5%) compared to when stimulation was OFF (M: 55.6%, SD: 4.5%) in the time window of the blink (Lag2 trials), but this seeming difference was reduced outside of the blink window, for Lag8 trials (ON-M: 68.9%, SD: 14.9%; OFF-M: 67.8%, SD: 3.4%). However, this effect did not seem to depend on distractor valence, as DBS similarly affected target accuracy in negative vs. neutral trials: In Lag2 trials ON stimulation improved performance for neutral distractors by 8.5% on average (SD: 13.6%, range: −6% to +26%), and for negative distractors by 7.1% (SD: 6.3%, range: +1% to +15%). This difference was either smaller or absent for Lag8 trials: accuracy improved by 3% for neutral (SD: 14.8%, range: −13% to +19%), and decreased by 1% for negative distractors (SD: 10.2%, range: −16% to +7%). In line with our expectations, DBS thus seemed to selectively improve accuracy on Lag2 trials, but contrary to our expectations, the valence or saliency of the distractor mattered little.

**Figure 3.**
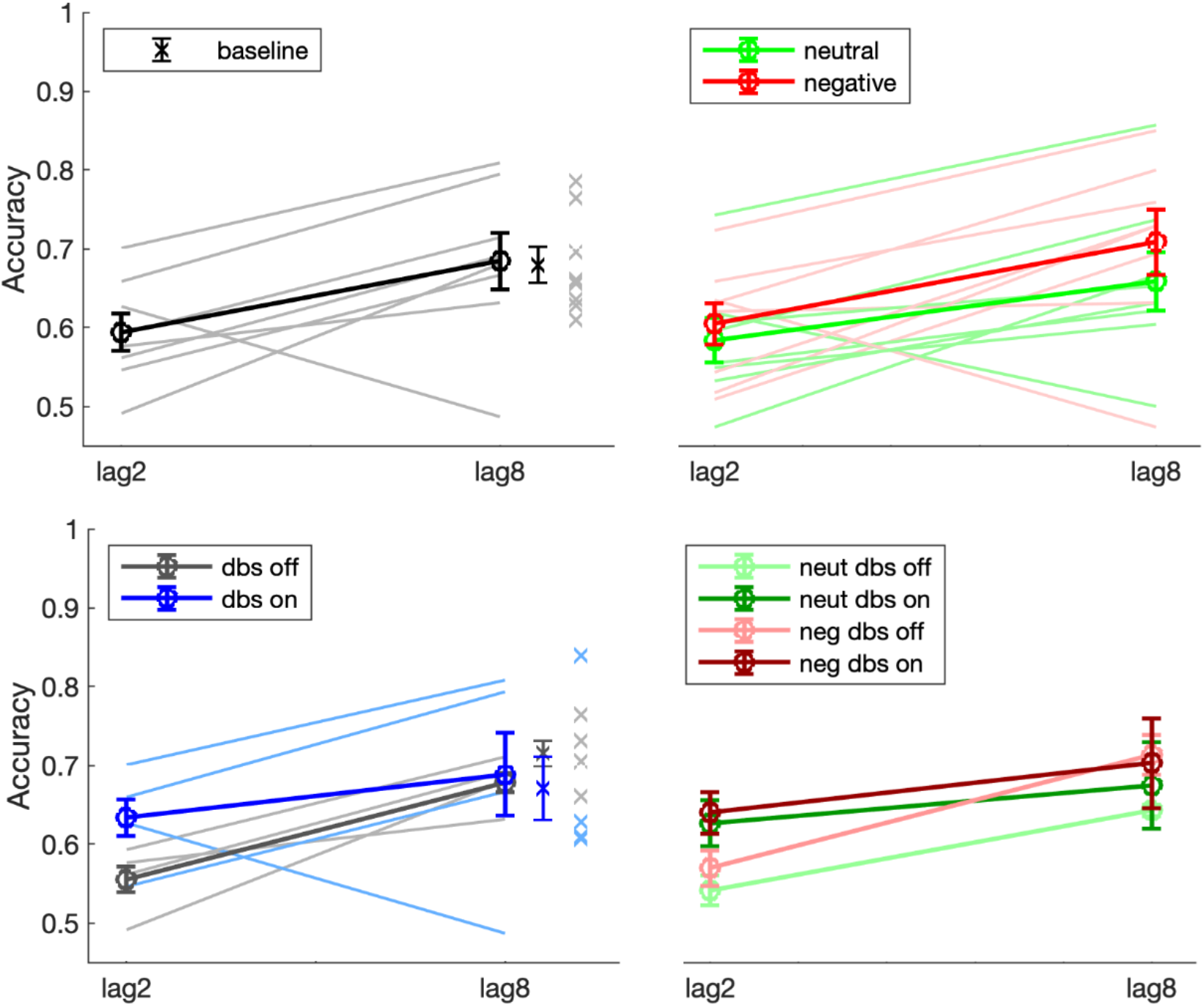
Mean accuracy for Lag across all eight sessions across four patients (top left), split by Valence (top right), DBS (bottom left), and their interaction (bottom right). Patients showcase a robust attentional blink on average, seemingly higher accuracy for negative distractors on Lag8 trials, and higher accuracy for DBS ON in Lag2 trials.

#### iEEG

We attempted to replicate and extend iEEG findings from Slagter et al (2017), where in a standard AB task, an increase in power in the α/β range (8-16 Hz) was present 80-140 ms after T1 (distractor in EIB) in T2 blink compared to no-blink trials, as well as an increase in the theta range (4-8 Hz) 150-400 ms after T2 was seen (target in EIB) in contrast with trials where it was unseen. Both expectations are marked in Fig. 4 with a black box. For neither the left or right striatal electrode no consistent effect can be seen after target presentation in the theta range in our data, neither OFF or ON DBS. There seems to be some consistency across patients for an effect in the α/β range on the left, but this effect was in the opposite direction from our prediction, and rather small in amplitude both for DBS OFF and ON, as can be seen in Figure 5. Thus, with our small number of cases, and a different blink paradigm, we did not reproduce the iEEG signatures of capture and conscious perception reported in Slagter et al. (2017).

**Figure 4.**
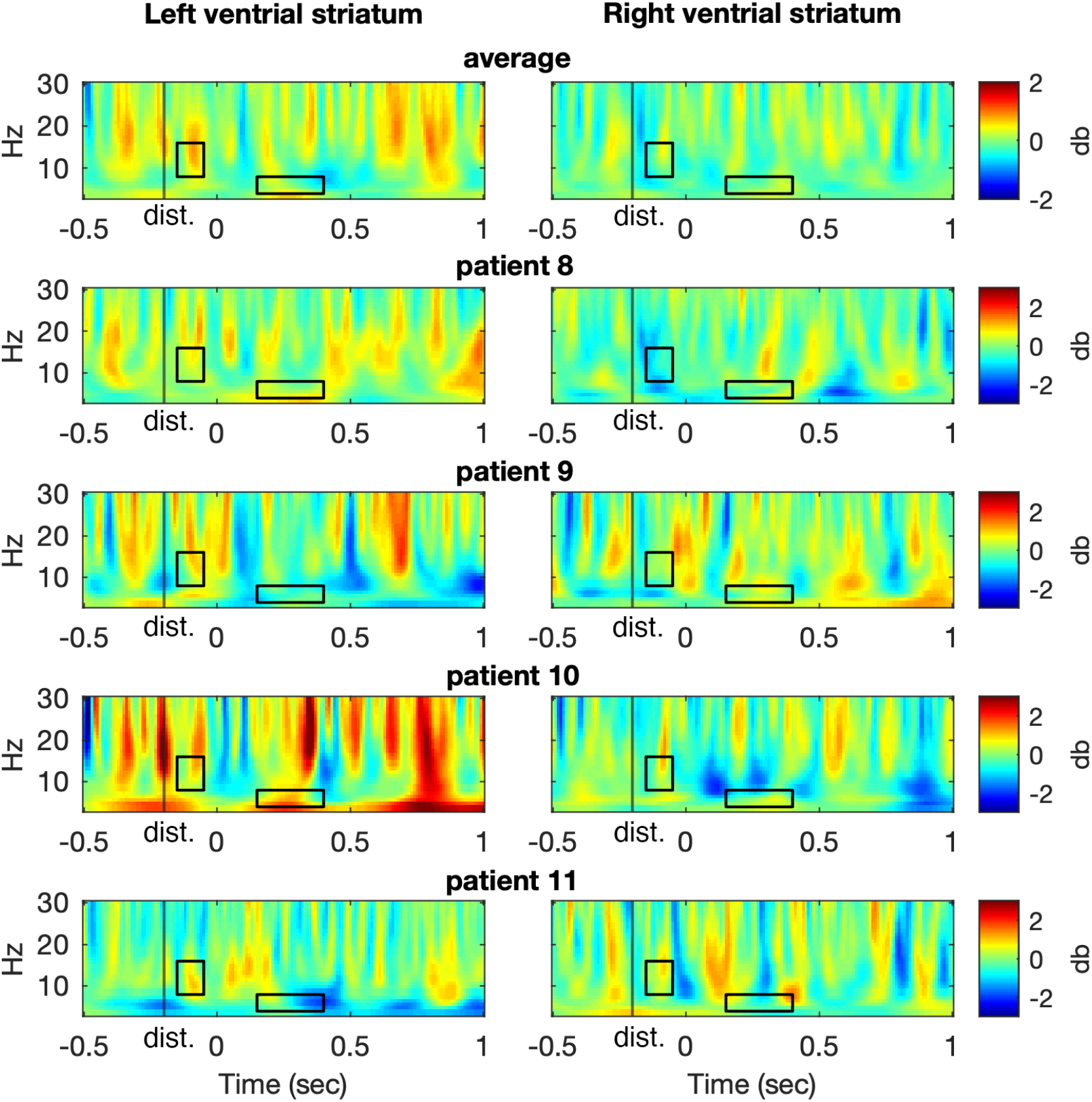
Time frequency results from the emotion-induced blindness task for seen versus unseen trials acrosss subjects (top row) and for each individual subject (lower rows). Presented are intracranial EEG data from the left (left figures) and right (right figures) ventral striatum. The target was presented at time zero. Time-frequency representations show in db the strength of differences in striatal activity between seen and unseen short-interval trials in the DBS OFF condition. The black rectangles denote the two time-frequency windows of interest.

**Figure 5.**
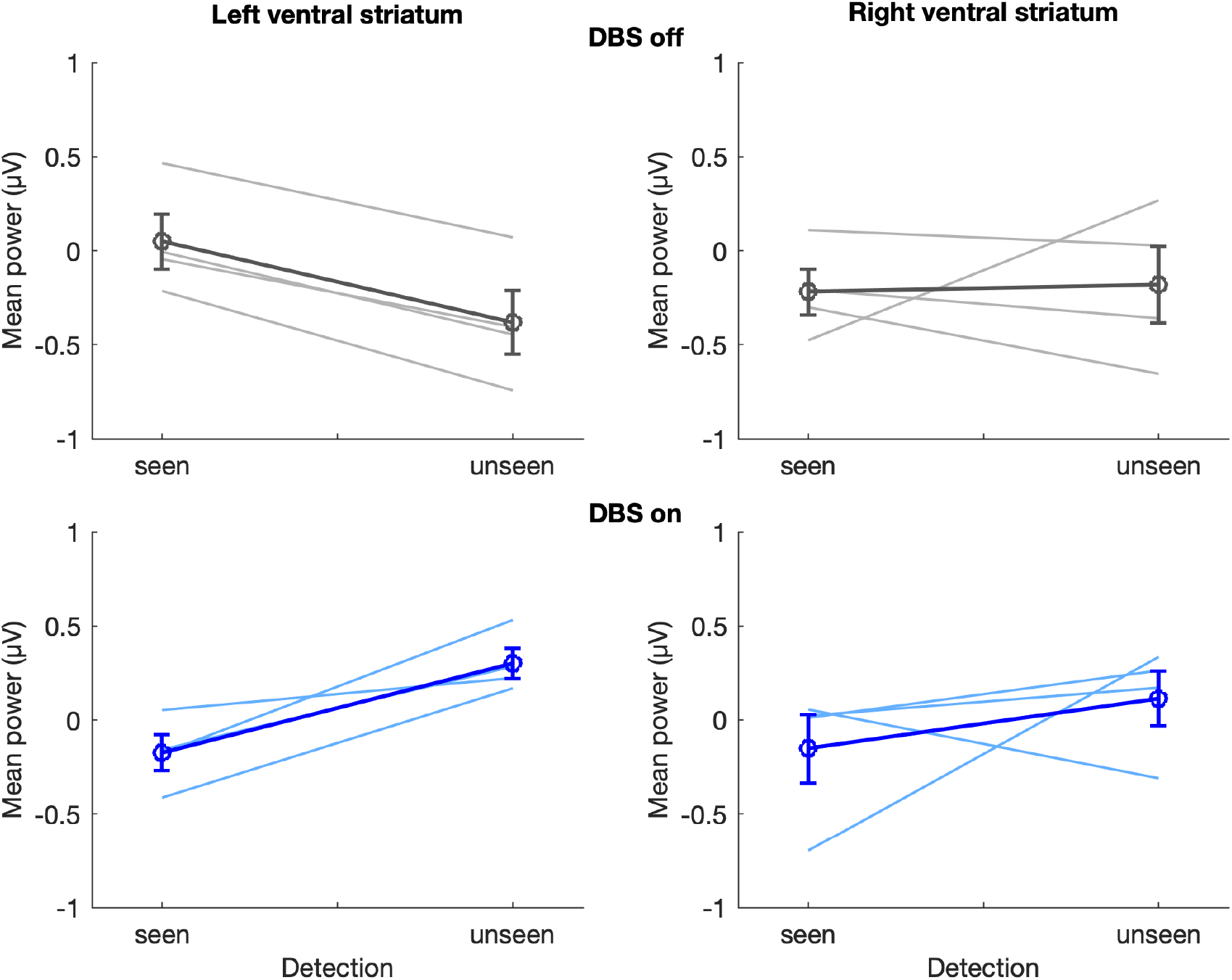
Presented is mean power for the predetermined α/β frequencies for the emotion-induced blindness task. As can be seen, we did not observe greater activity during DBS OFF in the α/β range to distractors in unseen vs. seen trials, in contrast to Slagter et al. (2017). For DBS ON as well, differences seem to deviate from zero only slightly.

### Backward masking

#### Behavior

As can be seen from Figure 6, patient’s accuracy improved with increased temporal asynchrony between target and mask (SOA), as is typically found (e.g., Breitmeyer, 2007). Whereas accuracy was at chance level for a SOA of 16 ms (M: 49.1%, SD: 5.8%), patients performed increasingly well with a SOA of 50 ms (M: 59.6%, SD: 10%) and even better at a SOA of 100 ms (M: 80.6%, SD: 17.4%). DBS ON improved accuracy for all three SOAs (16 ms M: +7.8%, SD: 5.9%, range: +2.5 to +14.2%; 50 ms M: +4.9%, SD: 8.4%, range: −1% to +14%; 100 ms M: +4%, SD: 10.9%, range: −2.2% to +16.6%). As this improvement was seen for all SOAs, and largest for the 16ms SOA, it is unclear if this effect reflects a true increase in accurate conscious perception or rather a response bias. This is unlikely however given we found little change in confidence scores for low SOAs (see below).

**Figure 6.**
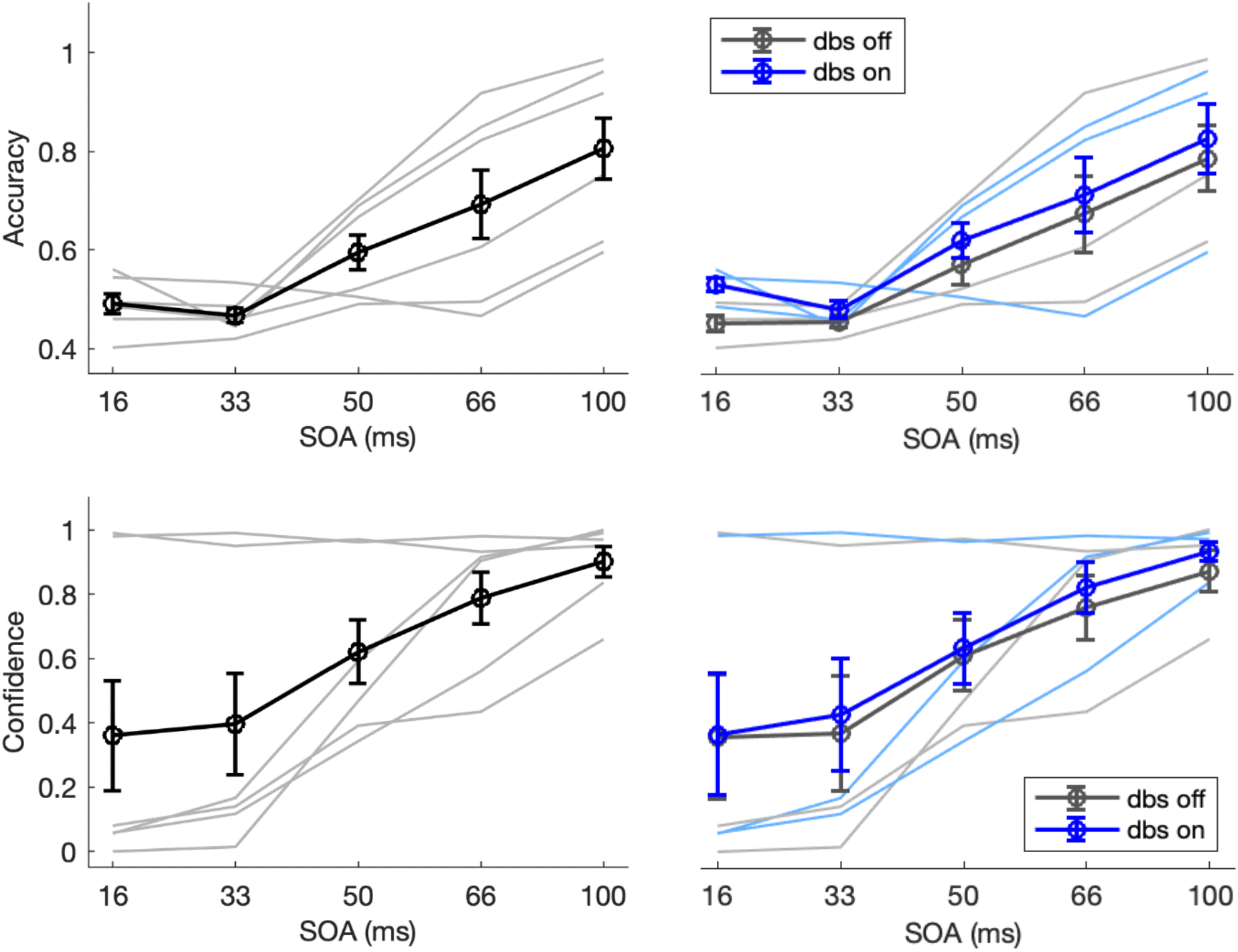
Mean accuracy and confidence, overall (left plots) and split by DBS conditions (right plots). Both accuracy and confidence scores improve with increased SOA (left plots; averaged across DBS OFF and ON). DBS generally enhanced accuracy, across SOA (top right plot), and confidence scores, in particular for the longer SOA durations (bottom right plot).

We observed a similar pattern for response confidence: patients were increasingly confident in their response with higher SOAs: 16 ms (M: 36%, SD: 48.4%) compared to 50 ms (M: 62%, SD: 28%) and 100 ms (M: 90.1%, SD: 13.2%). DBS ON also numerically enhanced confidence scores increasingly so for longer SOA durations: for 16 ms (M: +.8%, SD: 4.2%, range: −2.2% to +5.6%), 50 ms (M: +2.3%, SD: 9.1%, range: −4.8% to +12.5%) or 100 ms SOA trials (M: +6.2%, SD: 10%, range: −1% to +17.7%).

#### EEG

As shown in Fig. 7, we replicated previous reports showing that the target-evoked P3 ERP components are affected by the duration of target-mask SOA despite our small sample size (Boonstra et al., 2020; Del Cul et al., 2007). Specifically, the target-evoked P3 was virtually absent for the 16-ms SOA, in which targets were typically not seen, but present in all other SOA trials. While the amplitude of the P3 did not further scale with longer target-mask SOAs, as in our previous study (Boonstra et al., 2020), P3 latency seemed to scale with SOA. This could potentially be explained by a more pronounced preceding N2 in longer SOA trials (see Fig. 7), which may have temporally overlapped with the P3. Past work has shown that the N2 is the first ERP component to linearly scale in amplitude with conscious visibility (Sergent et al., 2005). A roughly similar pattern of findings could be observed for DBS ON.

**Figure 7.**
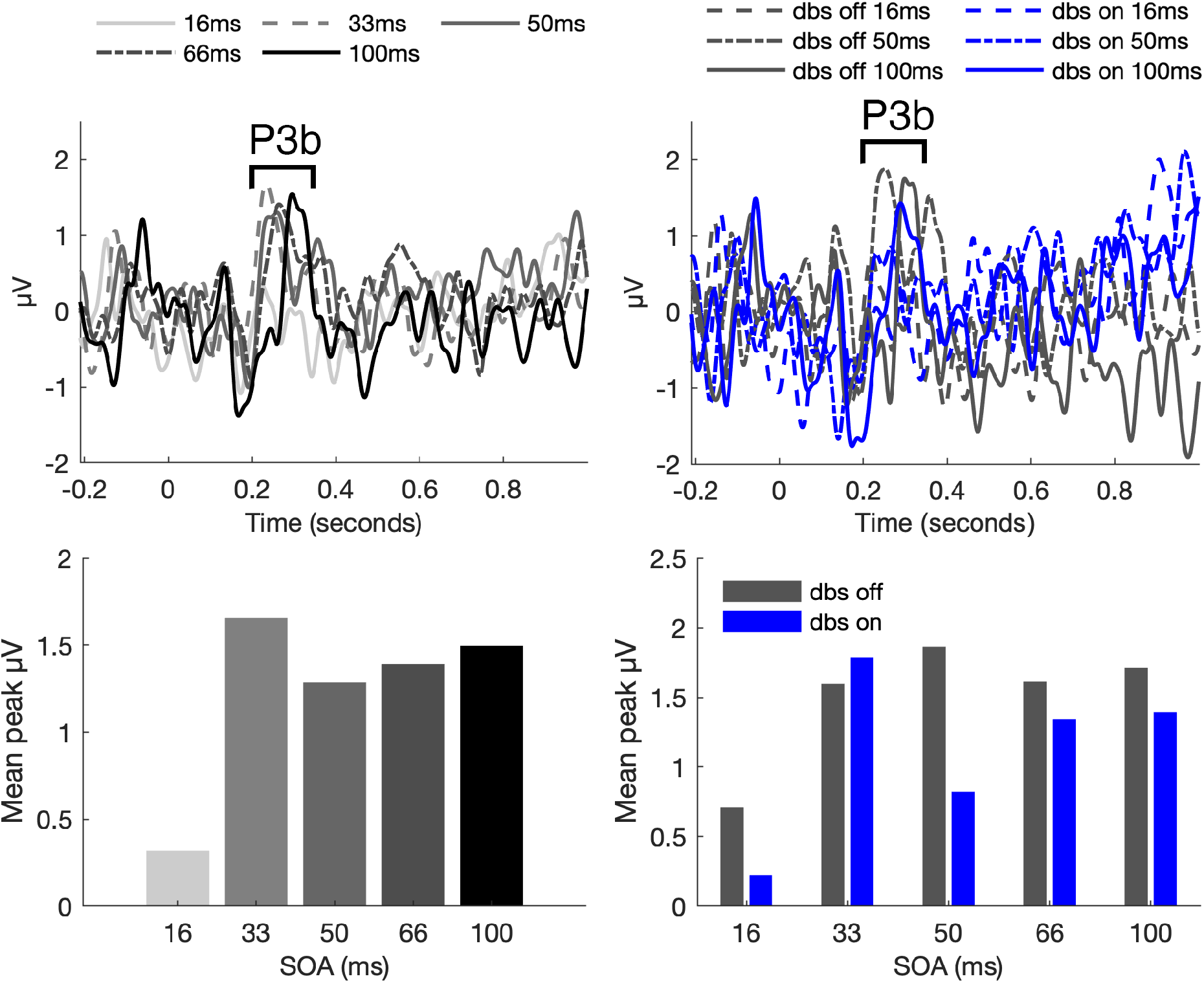
Effects of SOA and DBS on target-evoked ERPs in the backward masking task. This figure displays the grand-average target evoked ERPs for P3b electrodes (P1, Pz, P2, PO3, POz, PO4) for both DBS conditions combined (left panels) and separately (right panels), per target-mask SOA condition (16, 33, 50, 66, and 100 ms). This figure shows that ERP P3b amplitudes and latencies generally increased as a function of target-mask SOA, although not necessarily linearly (bottom figures).

#### iEEG

As in the case of the EIB task, we first tried to replicate the finding from Slagter et al (2017) concerning a selective theta response (4-8 Hz) between 150-400 ms to consciously perceived targets. As can be seen from Fig. 8, no such effect can be shown consistently across participants. Due to a lack of the expected effect in the OFF condition, time-frequency spectra for the ON condition are omitted.

**Figure 8.**
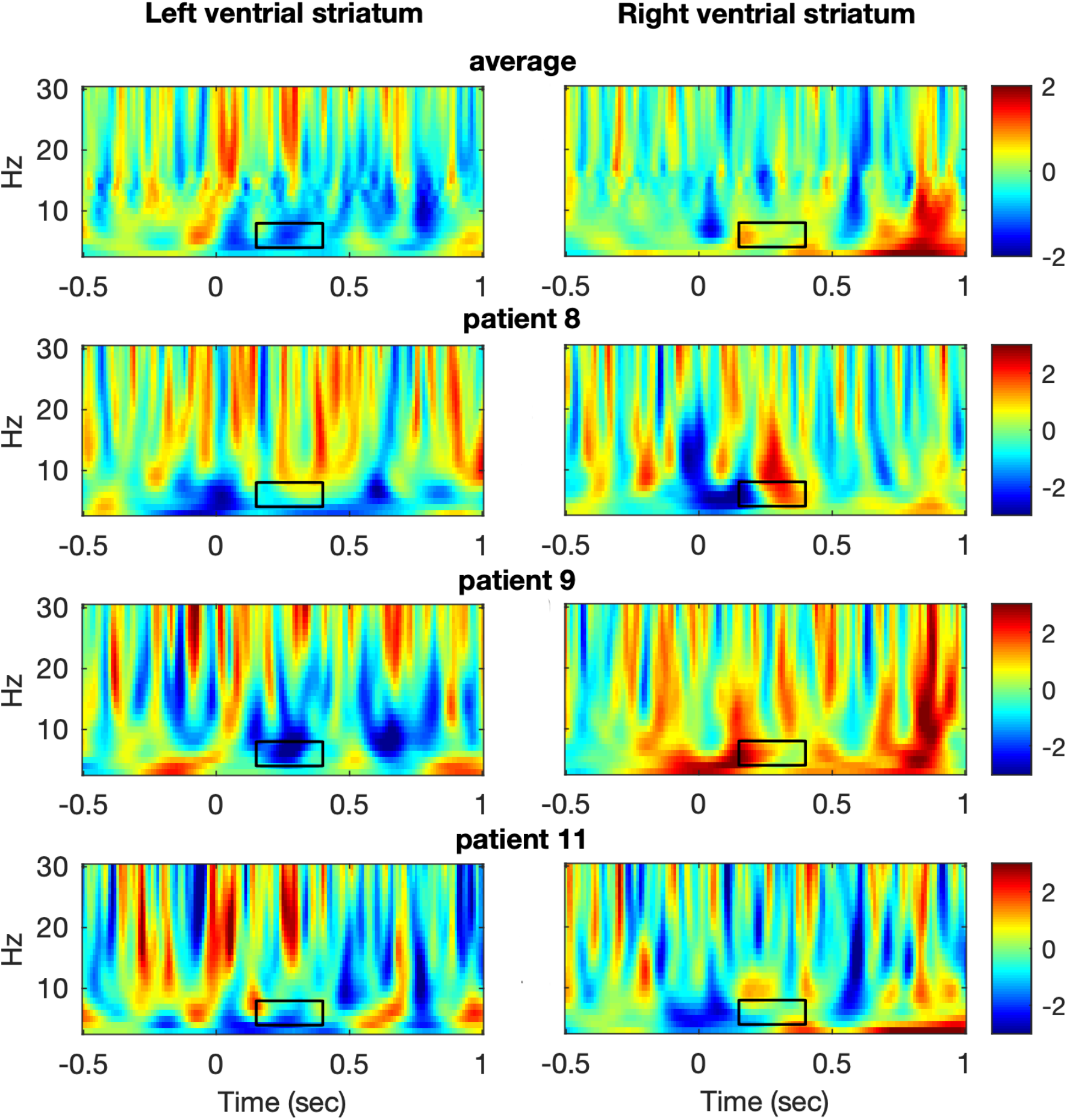
Time frequency results for seen versus unseen trials in the backward masking task OFF DBS. Intracranial EEG data are presented from the left (left figures) and right (right figures) ventral striatum. The target was presented at time zero. Time-frequency representations show in db the strength of differences in striatal activity between seen and unseen targets for 50 ms SOA trials with DBS OFF.

## Discussion

In this preliminary study, we presented results showcasing how deep brain stimulation in the basal ganglia may influence conscious perception. As a minimum requirement for the validity of our study, we showed how patients produced patterns of behavior expected for both emotion-induced blindness (EIB) and backward masking (BM). In addition, we showed that at a behavioral level there are indications suggesting that stimulation may improve target perception accuracy to a small degree in both tasks. Specifically, the emotion-induced blink was smaller ON DBS, and participants generally perceived more masked targets. Yet, our neural findings were less clear, conceivably due to our small sample size and their greater sensitivity to noise. We were able to largely replicate ERP findings from previous research for the BM task, showing a general increase in N2 and P3b amplitude and/or latency to consciously perceived targets as a function of SOA. Yet, we were unable to obtain reliable ERP responses to targets presented in an RSVP stream for the EIB task. Nor were we able to replicate time-frequency findings from previous research for either task OFF DBS. Lastly, we failed to show a meaningful effect of DBS on either ERPs or time-frequency decompositions. Thus, our findings suggest that ventral striatal DBS may improve conscious perception, but this critically awaits replication in large sample size studies that can also reveal the underlying neural mechanisms.

Currently, DBS is thought to affect neural tissue by generating efferent output at the stimulation frequency in the axon (Brocker & Grill, 2013; McIntyre et al., 2004). In the case of stimulation of the striatum in OCD patients specifically, the therapeutic effect of DBS seems to stem from the normalization of nucleus accumbens hypoactivity, and hyperconnectivity in the frontostriatal network (Acevedo et al., 2021; Figee et al., 2013; Smolders et al., 2013). More recently, two bundles of the anterior limb of the internal capsule have come to the forefront as target for stimulation: the anterior thalamic radiation (ATR) and the superolateral branch of the medial forebrain bundle (slMFB) (Coenen et al., 2012). The slMFB connects the prefrontal cortex via the nucleus accumbens to the ventral tegmental area (VTA), and contains dopaminergic projections from the VTA to the ventral striatum (Haber & McFarland, 1999). Improvements in mood are thought to contribute to the improvement of OCD symptoms through stimulation (Coenen et al., 2017). The ATR connects the prefrontal cortex to the anterior thalamus and is part of the cortico-striatal-thalamo-cortical network. Stimulation may normalize the dysregulation of this network in OCD (van den Heuvel et al., 2016). Stimulation of the slMFB seems most effective in combating OCD symptoms (Liebrand et al., 2019).

Given these courses of action, it is conceivable that the marginal behavioral improvements we recorded, stem from a normalized capacity for broadcasting of task-relevant stimuli, in line with theoretical accounts suggesting that conscious perception depends information being made globally available (Crick & Koch, 2003; Dehaene & Changeux, 2011; Edelman, 2003). Indeed, an explanation of our behavioral findings in these terms would support the proposed role played by the BG in perceptual processes (Redgrave et al., 1999).

It should be noted however that several limitations pertain to our study. We tried to replicate findings from a previous iEEG study from our group (Slagter et al., 2017), showing how a distractor (T1) elicited α/early β response, possibly reflecting attentional capture, was associated with failed perception of a subsequent target (T2), as well as a theta burst only for seen targets (T2). A direct comparison between this study and the present study is impeded by the fact that both tasks weren’t identical. While in the study by Slagter and colleagues (2017) a standard attentional blink task was used, in our study, patients performed an EIB task containing valenced distractors (equivalent to T1 in the AB task), and patients did not have to report the distractor. It must be emphasized there was good reason for this difference in design: we introduced a valenced distractor to strengthen the presumed attentional capture effect found previously with a neutral symbol. The idea was that if such a symbol was able to do so, then a negative image should elicit this capture effect to a greater degree, and we were interested to see if DBS could reduce this response.

Our sample size was reduced due to unforeseen limitations in DBS battery life, making it difficult to tell whether we succeeded in distilling signal from noise in our neural data. We also used a different stimulator than the study by Slagter and colleagues (2017). The unique advantage of this device is that it can stimulate and measure simultaneously, but a relative drawback is that the recording channel is online referenced to the top contact point on the same electrode through subtraction, which rendered it impossible for us to use the exact same referencing scheme as in Slagter al. (2017). Different contact points can show disparate and even opposite levels of activity, which complicates the comparison of our results with previous findings.

Despite these limitations, to our knowledge this is the first study to present data from patients performing an EIB and BM task ON and OFF DBS. We have shown there are indications to believe striatal DBS may improve conscious perception, but more research is needed to substantiate this claim.

## Notes

### Competing Interest Statement

The authors have declared no competing interest.

